# Spiking neural networks provide accurate, efficient and robust models for whisker stimulus classification and allow for inter-individual generalization

**DOI:** 10.1101/2023.04.19.537473

**Authors:** Steffen Albrecht, Jens R. Vandevelde, Edoardo Vecchi, Gabriele Berra, Davide Bassetti, Maik C. Stüttgen, Heiko J. Luhmann, Illia Horenko

## Abstract

With the help of high-performance computing, we benchmarked a selection of machine learning classification algorithms on the tasks of whisker stimulus detection, stimulus classification and behavior prediction based on electrophysiological recordings of layer-resolved local field potentials from the barrel cortex of awake mice. Machine learning models capable of accurately analyzing and interpreting the neuronal activity of awake animals during a behavioral experiment are promising for neural prostheses aimed at restoring a certain functionality of the brain for patients suffering from a severe brain injury. The liquid state machine, a highly efficient spiking neural network classifier that was designed for implementation on neuromorphic hardware, achieved the same level of accuracy compared to the other classifiers included in our benchmark study. Based on application scenarios related to the barrel cortex and relevant for neuroprosthetics, we show that the liquid state machine is able to find patterns in the recordings that are not only highly predictive but, more importantly, generalizable to data from individuals not used in the model training process. The generalizability of such models makes it possible to train a model on data obtained from one or more individuals without any brain lesion and transfer this model to a prosthesis required by the patient.

**Author Summary:** A neural prosthesis is a computationally driven device that restores the functionality of a damaged brain region for locked-in patients suffering from the aftereffects of a brain injury or severe stroke. As such devices are chronically implanted, they rely on small, low-powered microchips with limited computational resources. Based on recordings describing the neural activity of awake mice, we show that spiking neural networks, which are especially designed for microchips, are able to provide accurate classification models in application scenarios relevant in neuroprosthetics. Furthermore, models were generalizable across mice, corroborating that it will be possible to train a model on recordings from healthy individuals and transfer it to the patient’s prosthesis.

## Introduction

The development and improvement of brain-machine interfaces (BMI) strongly advanced during the last decades to process brain signals for the control of an external computer or machine [1, 2]. BMIs help paralyzed patients suffering from severe brain damage or disrupted connectivity between different brain regions, as typically caused by e.g. spinal cord damage, stroke or neurodegenerative diseases such as amyotrophic lateral sclerosis [3]. BMIs can read out and interpret brain activity to control external actuators such as robotic arms or to function as a spelling interface, thus enabling patients to write text and communicate with their social environment [4, 5].

Instead of turning neural activity into an action performed by a computer or machine, the interface can also be used to create a signal to be sent back to the brain. This would result in a closed-loop system that creates an artificial stimulation for the brain based on the signal it receives from it. Such systems are called brain-machine-brain interfaces (BMBIs) and are promising for neural prosthesis replacing impaired or even missing biological functionality [6]. For instance, it has been shown that such closed-loop systems can be used to bridge damaged neural pathways in the rat brain after traumatic brain injury by interpreting action potentials captured by multielectrode arrays (MEA) implanted in the cerebral cortex [7].

MEAs present several advantages, when considered for a neural prosthesis, as the shape of the MEA probe can be tailored to each individual case, flexible enough to adapt for different brain regions. On a high temporal resolution, they provide electrophysiological recordings which can then be further processed in different ways. One option is to use the local field potential (LFP), usually down-sampled and low-pass filtered to remove frequency components above 300 Hz [8]. The LFP reflects the gross spatially weighted average of membrane potential fluctuations of thousands of neurons within a few hundred microns around the electrode [9]. Another option is to extract action potentials or spikes from the high-frequency signal, usually band-pass filtered between 300 and 3,000 Hz. In the post-processing of the signal (spike sorting), spikes are categorized as single unit if the waveform can be clearly identified as action potentials of a single neuron. Otherwise, if spikes are overlapping, they are accumulated as linear combinations of action potentials from small neuron populations in the vicinity of the recording electrode, called multi-unit-activity (MUA) [10]. In both cases, computational approaches, implemented on the neural prosthesis, are challenged to accurately interpret the incoming data. An additional requirement is that these approaches must operate on small-scale and low-powered hardware suitable for chronic implantation but providing limited computational resources.

To explore the capability of computational methods able to comply with these constraints in a BMBI scenario, the rodent barrel cortex provides an advantageous experimental model [11]. Its prominent organization in cortical columns provides a one-to-one topographic representation of single whiskers, sensory organs on the animaĺs snout which can be mechanically stimulated in a well-controlled manner [12]. Whisker stimulation evokes a localized response in the barrel cortex that can be captured with MEAs and analyzed with the appropriate computational approaches like machine learning (ML) classification algorithms. Based on such recordings, it has been shown that ML classification algorithms, or classifiers, can accurately identify the cortical depth of the recording electrode and the deflection intensity of the whisker stimulus [13, 14]. While these authors could show that a pre-trained model can be applied on a small microchip within a reasonable amount of time, they did not provide an investigation of the capability of such hardware for training an accurate ML model online. This challenge was recently addressed by Petschenig *et al*. [8], who investigated and benchmarked ML algorithms for the classification of whisker deflection amplitudes based on MEA recordings from the barrel cortex of an anesthetized rat. Among the benchmarked methods, the liquid state machine (LSM) algorithm turned out to be highly accurate and this is particularly relevant since it implements a spiking neural network (SNN) specifically developed for neuromorphic hardware. Hardware of this type are small, brain-inspired computing architectures designed for processing brain signals in a highly efficient manner, representing an appropriate option for chronically implanted prostheses [15, 16].

The barrel cortex has therefore served as an ideal experimental model for investigating which computational approaches are appropriate for a neural prosthesis. The above summarized developments on the classification of the whisker stimulus intensity based on stimulus-evoked activity from the barrel cortex are highly promising for the field of neuroprosthetics, as they demonstrate the ability to implement BMIs or BMBIs in low-powered hardware and their capability to decode information from neural activity. However, these results are based on recordings under anesthesia, which is known to affect the pattern of neural activity [17]. Thus, it remains unclear if ML classifiers perform well also on data recorded from awake animals. Such data is expected to contain both a higher level of and a higher variability in spontaneous activity, compared to the anesthetized state, and therefore it could be more challenging for the algorithms to detect or classify stimulus-evoked activity. Moreover, it is not known if such models can be transferred from one individual to another. Such generalizable models would allow for training models on larger datasets composed of recordings from more than one individual, making the models more robust.

## Results

### Dataset and machine learning analysis

In this study, we analyze a dataset consisting of 19 MEA recording sessions and thus providing a large number of trials used to compose different datasets and consequently increasing the robustness of the models derived from these data. The data were obtained from the barrel cortex of four head-fixed water-restrained mice while they performed a go/no-go whisker stimulus detection task. Mice were conditioned to respond to whisker stimulation by licking at a water spout (**Fig. 1A**). During such an experimental session, a silicon probe with two shanks, 32 electrodes each, was used to derive electrophysiological recordings from the cortical layers (L) 2/3, 4, 5 and 6, which allows us to investigate which layer provides the highest predictive information within our ML-based analysis. Based on a current-source-density (CSD) analysis, we manually picked one electrode per layer to derive a layer-specific signal from the MEA recordings (see Methods).

**Figure 1.**
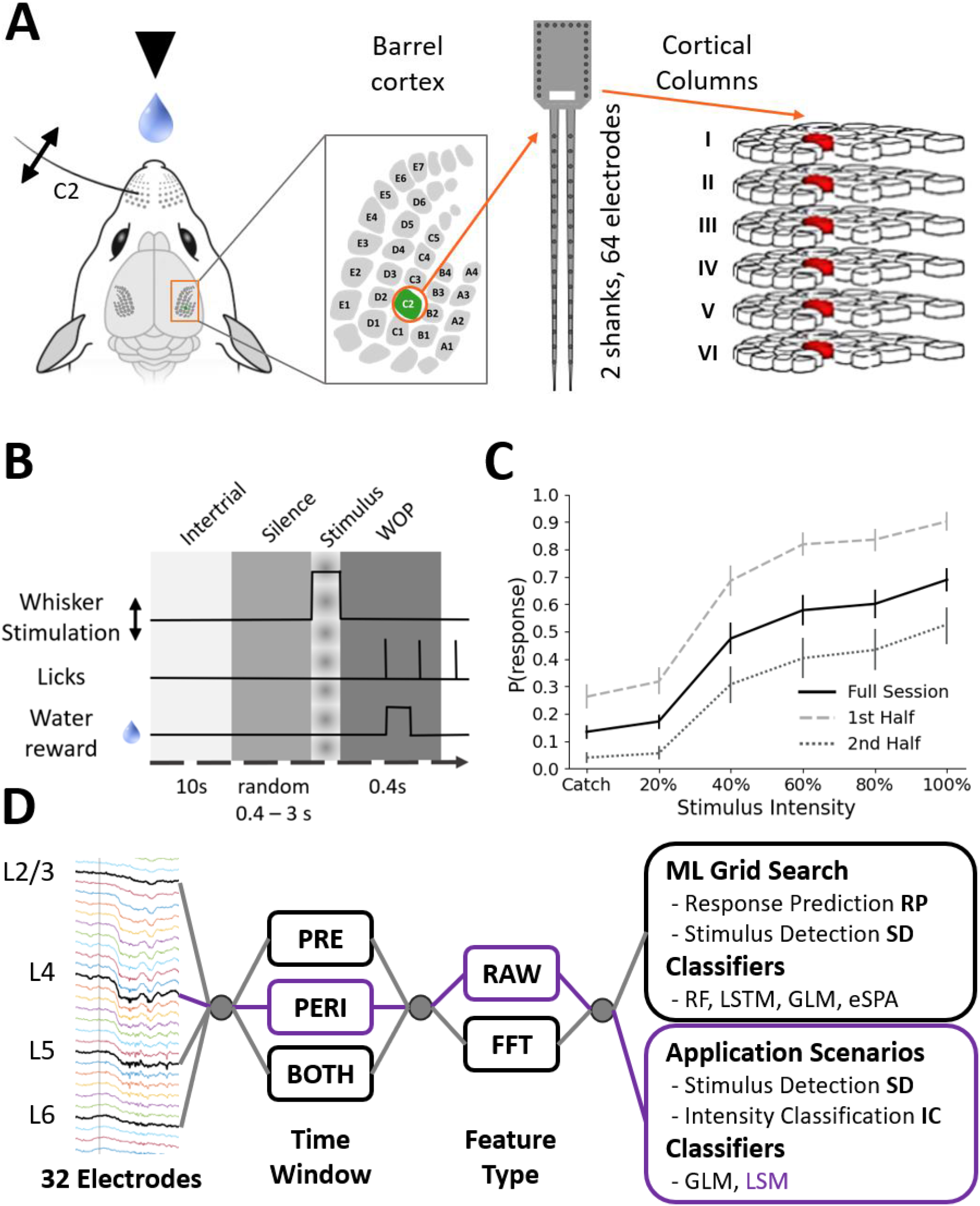
Experimental Setup and Psychometric Curves. **A** Head-fixed mouse within reach of a water spout. Multi-electrode-array recordings are derived from the cortical column associated with the stimulated whisker (here C2). A silicon probe with 64 electrodes, 32 on each of two shanks, was used to measure the activity from all cortical layers. **B** Overview of the behavioral task. Mice were operantly conditioned to respond to whisker stimulation by licking at the water spout. Responding on stimulus trials (but not catch trials) within 500 ms was rewarded with a drop of water. We refer to the time window from ∂400 ms to 0, relative to stimulus onset, as the pre-stimulus, or spontaneous, activity. The peri-stimulus or evoked activity is represented by the time window 0 to +100 ms as the stimulus is a sinusoidal whisker vibration of 100 ms duration. A *hit trial*, stimulus trial with a successful response of the mouse, is defined as a lick within the window of opportunity (WOP) from +100 to +500 ms. Trials with licks between 0 and +100 ms were excluded as licking activity caused strong artifacts in the LFP. **C** Psychometric curve averaged across 19 sessions from 4 mice. Only trials are considered that remain after all filtering criteria are applied (see Methods). Whisker stimulus intensity ranged from 0% (no stimulus) over 20, 40, 60 and 80 to 100% intensity (percentage of maximum deflection amplitude). Trials without stimulus (0% intensity) are called *catch trials* and were included to estimate the response rates achieved by random licks. Licks within the WOP in catch trials are called *false alarms*. Error bars show the standard error of the mean (SEM) computed over sessions. **D** Data flow for the machine learning grid search and application scenarios. One shank was selected that provides four electrodes from which the LFP was derived from the different cortical layers. The predictive features were then specified by the time window and feature type. In total three classification scenarios were investigated with 5 different classifiers, whereas only the peri-stimulus RAW features from L4 were used for the application scenarios in which well performing GLM classifiers were compared to the LSM (highlighted in purple). Subpanels in **A** were modified from [59, 60].

Each trial of the behavioral task consisted of different epochs (**Fig. 1B**). Before whisker stimulation, the mice had to refrain from licking for at least 400 ms (silence period). We used the 400 ms before stimulus onset to characterize spontaneous activity in the barrel cortex. The stimulus itself consisted of a 60 Hz sinusoidal vibration of 100 ms duration and we used this per-stimulus time window to analyzed stimulus-evoked activity. If the mouse responded within 500 ms after stimulus onset, it was rewarded with a droplet of water. Each recording session consisted of 300 trials, including 50 catch-trials without stimulation (0% intensity to assess spontaneous licking) and stimulus trials with different intensities ranging from 20% to 100% of maximum amplitude. Each stimulus was presented 50 times, stimulus sequence was pseudorandomized. As expected [18–20], response rate monotonically increased with increasing stimulus intensity (**Fig. 1C**).

As we integrated data from different recording sessions, we could not use spiking activity as each session provides a different set of single- and multi-units which disenables the assembly of a common feature vector to describe trials across sessions, a prerequisite for the machine learning algorithms. Besides this it has been demonstrated on a single-session dataset that (i) LFP features are as informative as features derived from MUA and (ii) it may not even necessary to derive more sophisticated features from the LFP, such as the response peak amplitude or response onset latency, but using the raw signal provides sufficient information for the ML methods used [8]. In general, the LFP recording is more stable in comparison to action potentials and therefore it is recommended to use the LFP in the context of neuroprosthetics [21, 22]. Following these previous studies, we use the raw LFP traces on 1,000 Hz resolution, low-pass filtered at 150 Hz but without further feature engineering, referring to them in the following as *RAW* features. Additionally, we investigate *FFT* features, consisting of the frequency components we obtain by applying the Fast Fourier Transform to the filtered signal [23]. Including the FFT features was inspired by Sederberg and colleagues, who demonstrated that field potentials in the barrel cortex can be informative with respect to how whisker stimuli are perceived in the barrel cortex of awake mice [24].

In summary, we trained models for stimulus detection (SD), response prediction (RP) and intensity classification (IC), which are the main specifications for the classification scenarios we investigated within the grid search and the application scenarios (**Fig. 1D**). The grid search was first done to investigate if there is predictive information within the LFP features towards the detection of a stimulus and the animal’s response to a stimulus. For these tasks, we benchmarked a set of four different classifiers, namely random forest (RF) [25], long short-term memory (LSTM) [26], the logistic regression as a generalized linear model (GLM) [27] and the entropy-optimal scalable probability approximation (eSPA) [28]. The latter algorithm was of particular interest due to its good performance on learning problems pertaining to the small data regime, to which most of our datasets belong to as they have a comparatively large number of features in relation to the number of observations [29]. The grid search was performed to investigate if classifiers established in natural sciences can find predictive patterns in the data and to evaluate which algorithm performs best. In an additional analysis with application scenarios relevant for neuroprosthetics, we involved also the liquid state machine (LSM), a classifier less-established in scientific research studies than those deployed in the grid search, but specifically designed for neuromorphic hardware [30]. Due to limitations with respect to software installation on the high-performance cluster (HPC), the LSM was not included within the more comprehensive grid search. However, the LSM was involved in the benchmarking based on the application scenarios relevant for neuroprosthetics in which we compared the runtime and accuracy of the LSM and the GLM, the method which showed the best performance during the grid search. The application scenarios involve the whisker stimulus intensity classification (IC) and a more comprehensive analysis of the stimulus detection (SD) (**Fig. 1D**).

### Grid search and best performing models

The first analysis addresses the question to what extent ML classification algorithms can leverage information from LFP features to train models for response prediction (RP) or stimulus detection (SD). Furthermore, exploring the results from the grid search reveals which classifier trains the most accurate models (**Fig. 2A**). Classification models achieve moderate accuracy scores for predicting the response of mice (RP) when using the peri-stimulus activity. A higher level of accuracy can be observed for SD models, reaching up to 90% when trials with 0% (catch-trials) and maximum stimulus intensity 100% were contrasted (CLEAR). Features derived from cortical layer L4 are more accurate than those derived from other layers or the averaged features (AVG). Considering a feature vector combing all layers (ALL) just slightly improves the accuracy over the performance of L4 alone. As expected, the peri-stimulus activity, describing the stimulus-evoked response in the LFP, is more predictive for both types of models in comparison to the pre-stimulus activity. As expected, models trained on the pre-stimulus activity features are not predictive for SD; however, for RP the accuracy is slightly increased compared to the baseline accuracy of 50% expected from a random guess model. An accuracy of 50% can be considered as a random guess performance, since the data set contains an equal number of positive (presence of stimulus or response) and negative (absence of stimulus or response) labels, due to the down-sampling of the most represented class.

**Figure 2.**
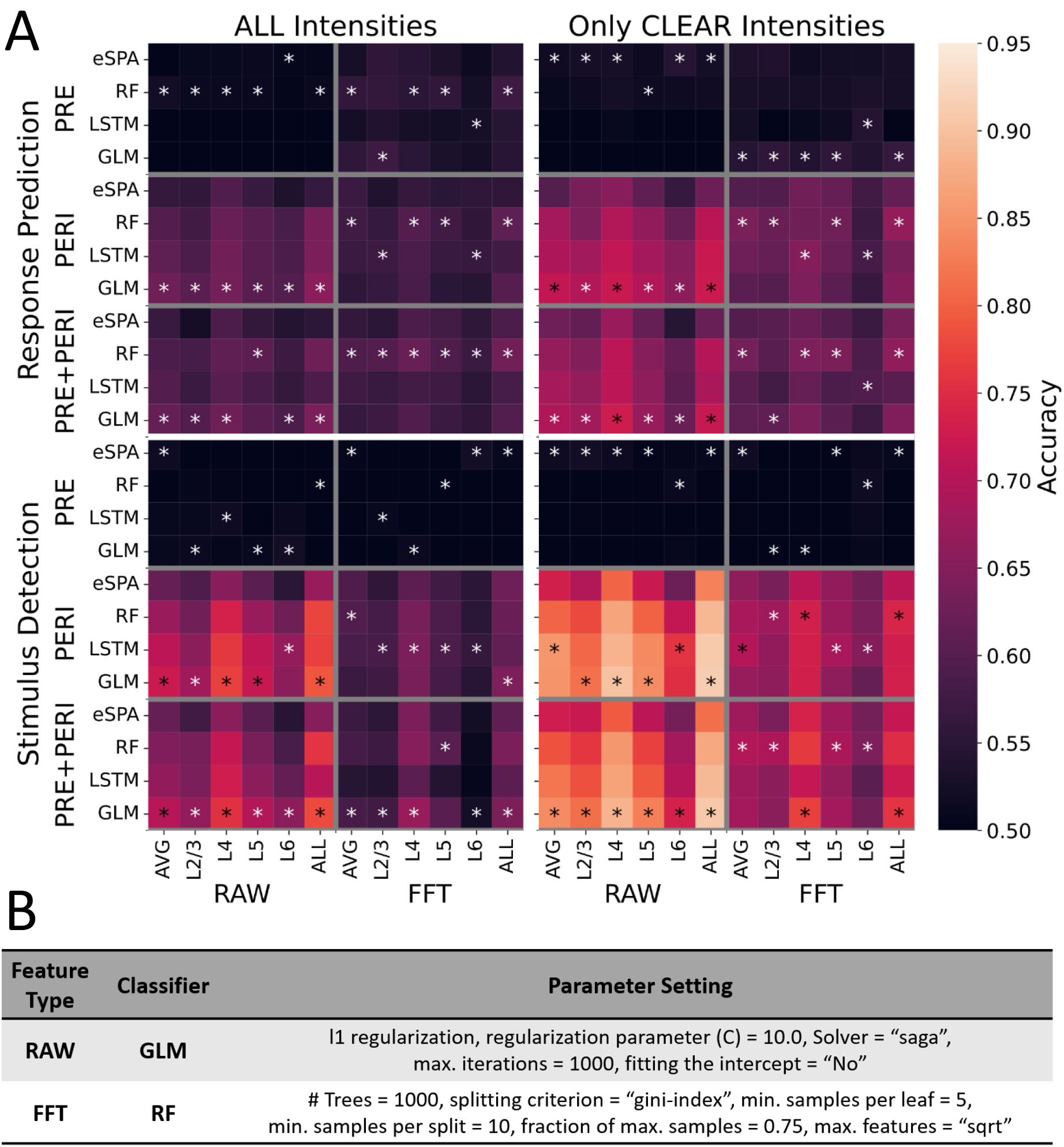
Machine Learning grid search and best performing models. **A** Classifier accuracy representing the predictive performance of four different classifiers in several supervised classification scenarios. These are specified by the time window before (PRE) or after (PERI) stimulus onset or the full time window (PRE+PERI). Furthermore, the feature types RAW and FFT define the scenario and from which layer the feature values were derived. The underlying datasets contain either trials from all intensities (ALL) or only catch trials (0%) and full intensity (100%) trials (CLEAR). The asterisks indicate which classifier achieved the best accuracy. **B** Details about those classifier settings that trained the best performing models for cases specified by the feature type.

Overall, we observed that the generalized linear models (GLM) with the underlying logit function (logistic regression) and Random Forests (RF) outperform the other classifiers. Furthermore, we found that the GLMs performed better when using the RAW features and RFs slightly improved the accuracy on scenarios with the FFT feature vectors. More precisely, based on the RAW features the GLM achieved the highest accuracy in 65% of the cases while Random Forest achieved the highest accuracy in only 45% of the FFT-feature scenarios. Based on this observation, we focused our attention on the two best performing models – indicated as the *generic best* models – which reached the highest validation accuracy averaged over all scenarios specified by the feature-type (**Fig. 2B**). In general, we do not see large differences between these selected *generic best* models and the *case best* models which are those with the highest validation accuracy for a specific scenario (**Fig. S1**). The only scenario in which we observed clear discrepancies between these two types of models is the one for SD detection from the ongoing activity, which has a predictive performance very close to a random guess model (**Fig. S1** – bottom panel, right-hand side).

A runtime comparison reveals that eSPA and the GLM require the lowest computational costs for training the classification model consistently across several scenarios (**Fig. S2**). LSTM is the most expensive algorithm while RF varies most with runtimes close to the cheapest classifiers for some cases but being sometimes as expensive as the LSTM in others.

In the following analyses, we focus on the best GLM setting and the RAW features, as we observed the highest accuracies for the corresponding models with the lowest training time. As mentioned, computational resources are limited in a neural prosthesis, and the runtime for model training usually depends on the number of features describing a trial. This holds true not only for training the model but also for evaluating a pre-trained model on new samples. Consequently, we proceed with the layer L4 feature set which is four times smaller than concatenated feature set (ALL) which provided only very small accuracy improvements over L4 alone.

### Conclusions on feature types and selection of relevant features

Before analyzing the RAW features most important for the generic best GLM model, we inspected the FFT features most important for the generic best RF model for stimulus detection (as RF performed best on the FFT feature). This inspection was done to better understand what information the FFT captures from the LFP, and why the models trained on the FFT features are worse than those trained on the RAW features. This analysis revealed that the RF model relies on the amplitude strength of the 30, 60 and 120 Hz frequency components (**Fig. S3**). Remember that the vibrotactile stimulus was a cosine of 60 Hz, moving the whisker in the anterior-posterior direction. Importantly, in the primary afferent neurons of the trigeminal ganglion (the first station of the ascending whisker sensory pathway), activity in response to whisker deflection is strongly determined by stimulus velocity [31]. This is also true for whisker-responsive neurons in the thalamus [32] and in barrel cortex [33]. For sinusoidal vibrations as we used, peak velocity is reached twice per stimulus cycle (one time in the anterior, one time in the posterior direction). Thus, oscillatory activity at two times the frequency of a vibrotactile stimulus is to be expected when having velocity-sensitive neurons. However, as shown by two example sessions (**Fig. 3A**), this oscillatory pattern is not always present, presumably because the recording electrodes have not always been in the close vicinity of the (supposedly small) neural populations that respond in that specific way. Thus, the FFT features describe a pattern that is not necessarily generalizable to other data as it is strongly induced by the shape of the external whisker stimulus, which is specific to our experimental setup.

**Figure 3.**
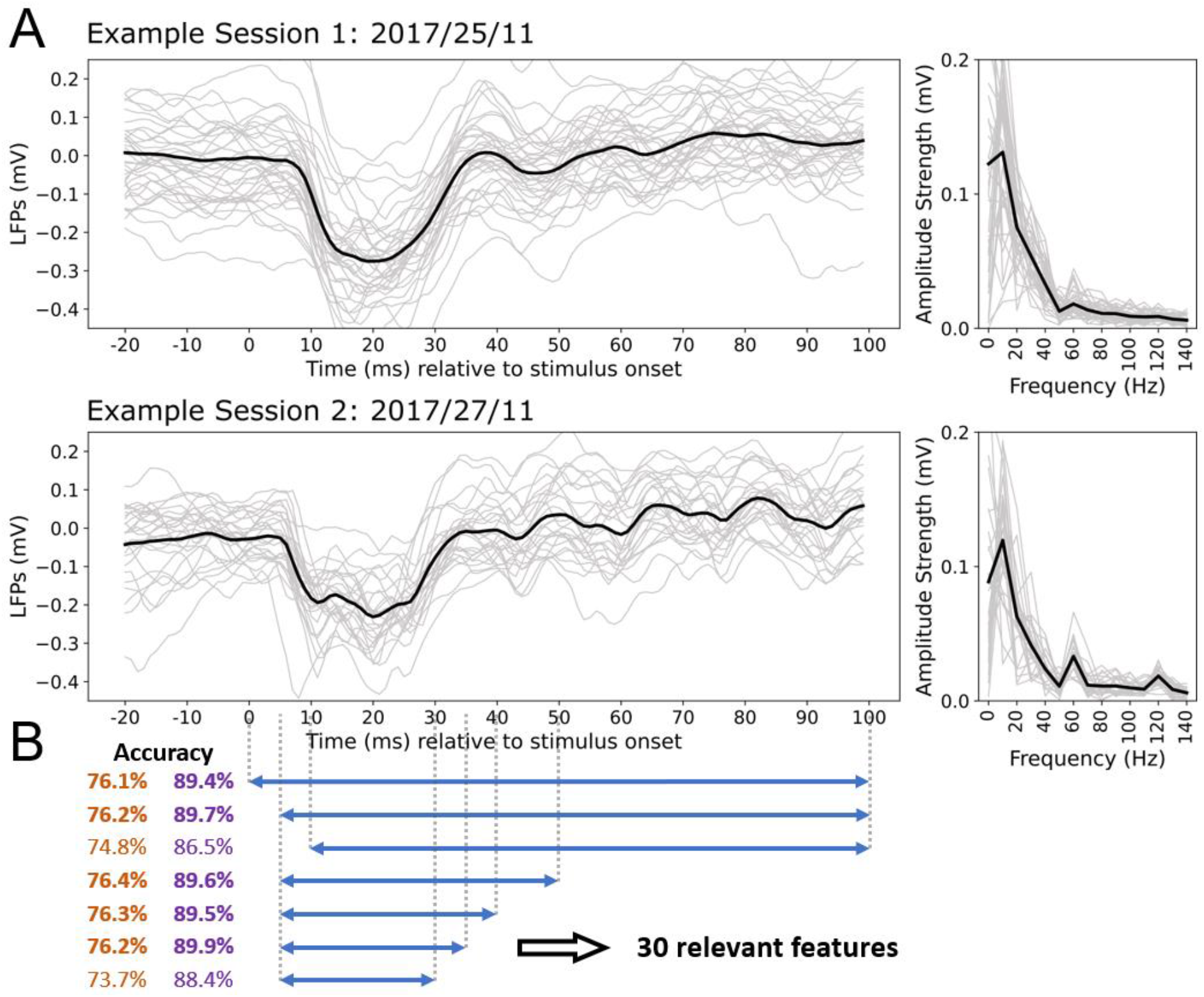
Selection of features, relevant for the ML model. **A** LFP recordings from two different sessions obtained with the L4 electrode and for 100% stimulus intensity trials only. Corresponding FFT spectra are shown in right panels. In the second example one can clearly see stronger amplitudes for the 60 and 120 Hz component while this oscillatory activity is not captured in example one. **B** Initially, we used the first 100 ms after stimulus onset in the grid search (upper, blue arrow). The GLM setting that performed best in the grid search was then used on smaller time windows (shorter arrows) and the accuracies were compared. Orange values describe the SD model accuracy on the data with all intensities while the purple values represent the SD model accuracy on only clear intensity trials (0% and 100%). Accuracy values that are similar to those achieved using the initial feature vector for the entire 100 ms are in bold. According to the accuracies achieved by the different models, the RAW trace from +5 to +35 ms provides the predictive information used by the classification model.

Those observations suggest that the RAW features offer a more solid representation, which could undergo a stronger feature reduction motivated by two arguments. The first is simply reducing the complexity of the ML task, as the runtime for training a model decreases when we decrease the number of features. The second, more important, reason is motivated by the observations we made on the FFT features, pointed out above and further described in the following. In both example sessions (**Fig. 3A**) we can see a strong, initial response to vibrotactile stimulation in the LFP, starting after ∼10 ms and lasting for approximately 20 to 30 ms. This agrees well with other studies investigating how the barrel cortex processes whisker stimuli [34, 35]. Only after this initial response, the recording might describe a 120 or 60 Hz oscillatory activity, as we can see in our data. However, the oscillatory activity is specific to the type of stimulus externally induced, and using the full peri-stimulus time window could potentially result in less generalizable models, biased by the stimulus shape. Using the best GLM setting we could eventually confirm that a smaller set of only 30 RAW features, describing the evoked response from +5 to +35 ms, contains enough information to achieve the same level of accuracy obtained by models relying on the complete pre-stimulus time window of 100 ms (**Fig. 3B**).

The grid search additionally revealed that models trained with the RAW features combined from all cortical layers did not perform much better than those trained on only L4 features. Again, we face a trade-off between accuracy and runtime [36]. As the complexity of the models increases with an increasing number of features, computational costs become higher not only for training but also for applying the model. In our case, these higher computational costs cannot be justified by minor improvements in accuracy. More precisely, while the model complexity increases by factor four when all layers are used compared to only L4 features, the GLM model for SD in the CLEAR scenario improved by only 1.3%.

Interestingly, the GLM training time decreases when all features are used compared to the single layer (L4) model in the scenario mentioned above (**Fig. S2**, two center panels). This could be explained by small changes in the parameter settings tuned within the grid search, but also by overhead presumably related to file-system operations on the HPC system on which the grid search was conducted. Hence, the grid search runtimes are useful to get an orientation, but not most accurate. For these reasons, the more precise runtime comparisons that involve the best generic GLM settings in comparison to the LSM algorithm (see next section), were done on a local machine on which we had full control over the resources used. Here we ensured that the computational resources available for the model training is the same for each algorithm and here we could also exclude any other factors that might influence the computational processes on the machine while performing the runtime comparisons.

### SNN benchmark on the neuroprosthetics application scenarios

After identifying the best classification algorithm and an optimal set of features for stimulus detection, we analyzed two application cases relevant for neural prostheses. These are the detection of stimuli of different intensities and, inspired by previous related studies, classification of stimulus intensity (IC). Both classification tasks represent biological functions covered by the barrel cortex, and having an accurate ML model for these tasks would indicate that a classifier could successfully be applied on a prosthesis restoring these functionalities. Being able to demonstrate that such models are highly accurate corroborates that ML can be applied for other brain regions as well, restoring other, possibly even higher-order, functions of the brain.

In the classification tasks, we also considered a particular type of SNN, the liquid state machine, as an algorithm specifically designed for the implementation on neuromorphic hardware [30, 37, 38]. In terms of predictive accuracy and model training time, this algorithm was compared to the generic *best GLM* and additionally to the *slim GLM* that has the same setting as the best GLM but using a lower number of maximum iterations (100 as the default). The slim GLM setting was involved to analyze if a shortened training results in lower model accuracy and to additionally challenge the supposedly short training time of the LSM. Compared to both variants of the GLM, the LSM achieves similar accuracies for all cases of SD and IC models (**Fig. 4A**). As expected, the stronger the stimulus intensity (SI) the more accurate is the detection, with very poor predictive performance for the lowest intensity. To evaluate the IC models, we created five classification scenarios, with trials from two intensities differing by 40% in the first three cases and differing by 60% in the remaining two cases. Even though the first three cases have the same difference in intensity, the model accuracy is higher for the first case of “20% vs. 60%”. The models, differentiating stimuli with 60% difference, tend to be more accurate. However, they can be on the same level as we can see for the example cases “20% vs. 60%” and “40% vs. 100%”.

**Figure 4.**
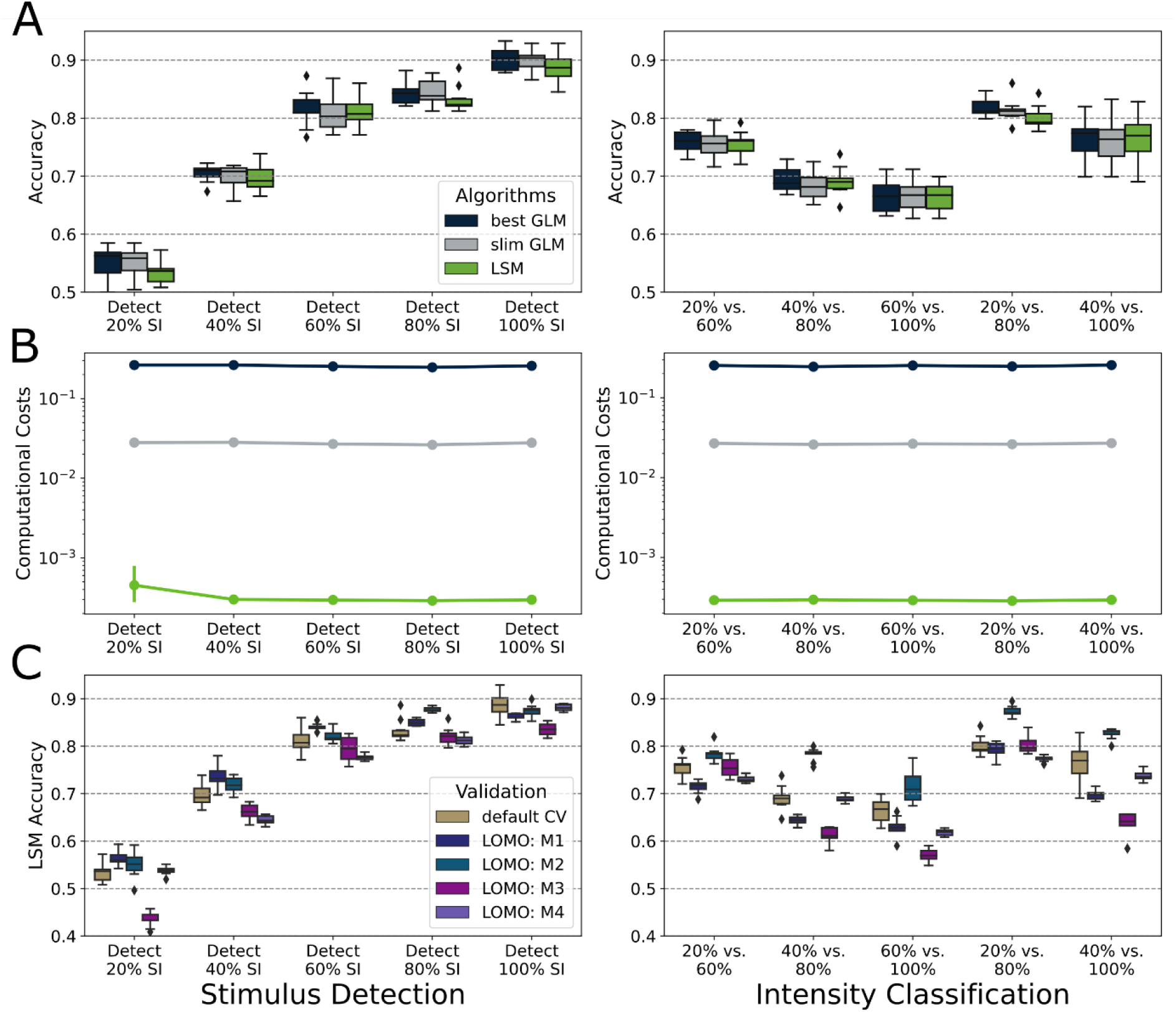
SNN benchmark in application scenarios. The application cases, relevant in neuroprosthetics, are whisker stimulus detection (left panels) and stimulus intensity classification (right panels). **A** Comparing the accuracies achieved by the *best GLM* setting defined during the grid search, another *slim GLM* with same setting, but applying a faster exit condition and the liquid state machine (LSM). **B** Same methods as in **A**, comparing the computational costs represented by the model training time in seconds. **C** Additional leave-one-mouse-out (LOMO) validation in comparison to the default cross-validation that includes a mixture of trials representing all individuals.

Compared to the best GLM, the slim GLM requires less computational costs, as expected. However, in terms of computational cost, it is also unable to outperform the LSM which decreases the runtime by two or three orders of magnitude compared to the best GLM or slim GLM, respectively (**Fig. 4B**).

Using the LSM we then investigated the performance of models when trained on trials excluding the data from one mouse that was left out from training but used to validate the model. This concept, called leave-one-mouse-out (LOMO), is analyzed in comparison to the cross-validation concept used so far (see Methods). The LOMO validation applies a bagging (bootstrap aggregation) concept to obtain more robust accuracy scores, a conventional cross-validation was not applied as we needed to separate trials based on the mouse used for the recording. For the higher intensities, with values above 40%, we do not see strong differences in performance when the model is applied to data from an individual completely excluded from the training process (**Fig. 4C**). This holds true for the IC models as well.

## Discussion

Inspired by brain-machine interfaces (BMIs), the brain-machine-brain interface (BMBI) is a key concept in neuroprosthetics [39, 40]. On a chronically implanted neural prosthesis, low-powered computer architectures, designed to accurately read out, analyze and interpret brain activity, are implemented to replace the functionality of a damaged brain region. Machine learning (ML) algorithms are promising to be applied on such hardware due to their high predictive accuracy on classifying evoked neural activity as has been shown on multi-electrode array (MEA) recordings from cerebral cortex of anesthetized animals [41–43]. Moreover, it is of paramount importance to employ algorithms with minimal computational cost, not only to achieve real-time data processing but also for efficient hardware usage. Spiking neural networks seem to be most efficient and provide accurate models [44], however, it has not been investigated how such classifiers perform on data from awake animals which is likely to be more challenging, for example because of fluctuations in neural activity related to sensory and cognitive processing. It has also been unclear if such models are generalizable from one individual to another, a potentially important aspect as this would allow for using a model trained on more comprehensive recordings from many, also healthy, individuals to be transferred to a neural prosthesis.

We benchmarked widely used ML classifiers on a comparatively large-scale electrophysiological dataset describing neural activity in the barrel cortex of four mice performing a go/no-go whisker stimulus detection task. The classification tasks we investigated include the detection of a whisker stimulus and the differentiation of stimulus intensities, both tasks related to functionalities of the barrel cortex and, considering the experimental model we use, relevant for a neural prosthesis. On these scenarios, we also investigated the accuracy and runtime of the liquid state machine (LSM), a spiking neural network especially designed for neuromorphic hardware and known to be very efficient in model training. Based on our data, we confirmed that the LSM is as accurate as the competing algorithms. Furthermore, we could show that the models allow for inter-individual generalization. Our findings are discussed in the following sections together with an outlook for future work.

### Algorithm comparison on stimulus detection and classification

Artificial neural networks, such as the LSTM, are promising for a large variety of applications, including challenges from neuroscience [45]. On our datasets, however, the LSTM was not only unable to outperform the other algorithms, but also consumed way more computational resources for the training process. In contrast, the generalized linear model turned out to be very accurate with comparatively low costs on model training. Eventually, on the neuroprosthetics application scenarios, we could see that using a spiking neural network could further and considerably reduce the training time as compared to the GLM, while achieving similar accuracies. Even though many of the datasets we analyzed are from the small-data learning regime, characterized by a strong imbalance between the small sample size and the comparatively high number of features, the eSPA algorithm was not able to perform as well as its competitors, which is unexpected due to its superior performance on other small-data learning challenges [28, 29]. We assume that this is explained by the nature of the data, as the shape and strength of a stimulus-evoked response in the barrel cortex varies strongly between trials even when considering trials from the same recording session and with same stimulus intensity (see examples in Fig. 3B). In a d-dimensional space, where *d* is the number of features, this leads to a continuous transition from catch trials to stimulus trials. The logit-function used within the GLM optimization is expected to represent such transitions way better than the discrete clustering approach implemented by eSPA. To overcome this limitation, the current version of eSPA could be extended by a soft-clustering approach as used by the initial SPA method [46]. This would be beneficial in the context of neuroprosthetics, because eSPA models are not only CPU- and memory-efficient, and therefore suitable for microchips, but they are also interpretable, in contrast to LSM models for which this aspect represents an open challenge. From this perspective, eSPA is advantageous, because interpretable BMIs can potentially reveal deeper insights into biological concepts of how neural information is processed [47].

We observed a high accuracy of the generalized linear models trained on the RAW features, especially for detection of the stimulus with the highest intensity. This strongly suggests that the RAW features do not encode complex, non-linear, but rather linear patterns mainly characterized by the strength of the response in the LFP. Otherwise, more complex classifiers as the RF, LSTM or eSPA would have shown superior performance in the grid search as these algorithms are able to leverage non-linearity in data. Considering this, it is even more important to explore algorithms that are as simple as possible when it comes to algorithmic complexity resulting in a short model training and little memory usage, but complex enough to achieve competitive accuracy. These aspects motivated us to additionally explore the potential of the LSM algorithm.

### Response prediction and the patterns in spontaneous activity

Besides investigating neural activity with respect to how whisker stimuli are processed in the barrel cortex, the dataset allowed us to search for patterns predictive towards the behavioral response. This is a critical aspect as we record from a somatosensory brain region believed to be involved in the process of evaluating an incoming stimulus and provide the basis for sensory-driven decision-making [12]. Considering the response prediction analysis as part of the ML grid search, we observe a moderate accuracy of around 72% for models trained on the RAW features of both pre- and peri-stimulus activity (**Fig. 2**). However, an additional analysis based on the predictions of the corresponding models, strongly suggests that the response prediction models basically perform stimulus detection, which results in quite accurate models due to the strong correlation between the presence and absence of a stimulus and the occurrence of a behavioral response (see **SI Results, Section 1, Fig. S4**).

Interestingly, our ML grid search revealed that patterns exist in the harmonic oscillations derived from spontaneous (pre-stimulus) activity that are slightly predictive towards the response of mice (see RP with pre-stimulus FFT features, ALL intensities, **Fig. 2**). For the FFT features derived from L2/3, L4 or L5 we observed accuracies slightly, but significantly better than the random baseline when all stimulus intensities are considered in the data (p-value from Fischer’s test < 1e^-26^ in all three cases). Even though this difference in accuracy is small, these results indicate that neuronal activity in the barrel cortex is also related to attention or anticipation of the expected whisker stimulus [48, 49]. Such weak patterns in the spontaneous activity could reflect brain states in which the mouse is more attentive to perceive the stimulus which has an impact on the response especially when not only the full intensity trials but also those with a small or intermediate intensity are analyzed.

### Efficient and accurate models for neuroprosthetics application scenarios

Regarding the application scenarios, we could show that the liquid state machine is a very promising approach which lends itself well to implementation on neuromorphic hardware. It achieves the same level of accuracy as the GLM, but requires significantly less computational resources. In agreement with the findings of Petschenig *et al.* and Wang *et al.*, the most informative signal is derived from layer L4, as we observed in the grid search. Previous in vivo recordings in rat barrel cortex have identified inhibitory interneurons in L4 as the units carrying the highest amount of sensory stimulus-related information [34, 50]. Thus, neuronal activity in L4 represents a good target to obtain information on the spatio-temporal properties of the sensory input. Measuring the activity from L4 is not only possible, but also very reliable with invasive technologies as used in our experimental model, however, in a practical application it will be more beneficial to use non-invasive technologies to avoid an intervention in the patient’s brain. Resolving activity in a specific portion of tissue without having direct access could be realized by laminar inference based in MEG recordings [51], although, MEG is not applicable in neuroprosthetics.

Considering the scenario of stimulus detection, the benchmarked algorithms achieve a high accuracy of ∼90% for the strongest intensity and the detection of 60% and 80% stimulus intensity trials was still possible with an accuracy greater than 80% (**Fig. 4A**). Thus, the LSM could accurately perform such a task on the microchip for different stimulus intensities. The ML models achieved a lower accuracy on the detection of stimuli of lower intensity (< 60%) which agrees with the lower response probability of mice as observed in the psychometric curve (**Fig. 1C**), suggesting that mice had more difficulties to perceive, in other words detect, such low-intensity stimuli. In comparison to the 100% stimulus intensity detection, the level of accuracy was also lower considering most of the SI scenarios (**Fig. 4A**), even though the well-calibrated ML algorithms were used. Thus, the LFP patterns that distinguish stimuli from different intensities are not strong enough to be accurately learned during the model training or the algorithms are not able to capture these patterns. However, before further investigating strategies to improve the accuracy of classifying such data it might be important to consider other types of data for future work. We suggest this as the datasets that have been used so far, including the one we used, are based on precisely controlled single-whisker stimuli, certainly different from how whiskers bent and move when animals are actively palpating during unrestrained exploration [8, 13, 14]. Therefore, for future work, it would be highly relevant to apply and test the suggested algorithms on recordings from microchips implanted in freely moving animals. The hardware, required to do this, could use wireless and battery-free devices [52, 53]. Recordings from such experiments might result in LFP traces that are less distinguishable as we would expect that whisker deflections from freely moving animals are less distinct. In such a context, it could be of interest to explore the potential of supervised machine learning based on regression models providing predictions for continuous labels in contrast to the discrete output from classification models.

To realize the closed-loop system, the output of the ML model can be used to control an electrode located in another brain region to induce intracortical microstimulation (ICMS) that can convey artificial signals that may be as complex as the somatic perception of tactile properties from different objects [54, 55]. Considering the flexibility of ICMS there should be no limitation regarding the artificial stimulation based on potentially comprehensive predictions from the ML model even if those are continuous labels as provided by regression models.

### Model robustness and generalizability

Another important aspect is the robustness of the findings concluded from our ML-based analyses. The sample size is a key factor in statistical analyses and ML models tend to be more meaningful and generalizable when trained on large datasets [56, 57]. Even though electrophysiological recordings are challenging from different perspectives, we could explore a comparatively large dataset using 4 selected electrodes, out of 64 and > 3,400 trials for 6 stimulus intensities and two types of behavioral response, go and no-go. The final subsets used for the ML-based analysis were restricted by a selection of trials for the different intensities resulting in a sample size of at least 1,000 trials, which is approximately four times higher than in comparable studies using only one recording session, e.g. [8, 13, 14]. Within the LOMO validation, the training set was always limited to data from three out of four mice and the size depended on the number of sessions available for the different animals. The minimum training set size was > 600 trials for these cases, including cases with > 1,000 trials as well.

Leveraging the big advantage of our data, which includes recordings from different individuals, we could show that recordings are highly stable across sessions. The liquid state machine performed well on the corresponding application scenarios. This holds true even when models are evaluated on data from individuals that were completely excluded from the training procedure (**Fig. 4C**). This strongly indicates that models trained on healthy neuronal activity can be transferred to a prosthesis, eventually used by patients.

In summary, based on a large electrophysiological dataset enabling a robust model evaluation, we could show that SNNs are accurate and efficient in interpreting neural activity towards neuroprosthetics application scenarios related to the barrel cortex. By using the LSM classifier, the model training could be realized on a small microchip to be implemented on the prosthesis which can be beneficial to retrain a model for adjusting to small drifts of the implanted electrode, for instance. Note that in all application scenarios the LSM was two orders of magnitudes faster than the optimized (slim) GLM and achieved a model training time of less than one millisecond (**Fig. 4B**). While our tests were done on a usual local machine with the main perspective to compare the model training time between the different classifiers, it has been shown recently that LSMs can efficiently be implemented and executed on microchips, demonstrated on a DYNAP-SE neuromorphic processor [8]. Another opportunity is to train a model on an external machine, transferring the model and only applying it on the prosthesis [14]. If future technologies enabled a more stable implantation of the electrodes, retraining on the prosthesis might not be necessary. Our generalizability analysis confirms that using a pre-trained model is possible even if the underlying data was derived from an individual different from the patient who needs the prosthesis. Hence, it is not even necessary to derive many recording samples from the patient to receive a large dataset for training a robust model, it would be sufficient to just capture the neural activity as input for the prosthesis that implements a model, already trained externally on a large dataset. As the LSM was the fastest algorithm, we report its performance in the generalizability validation (LOMO) in **Figure 4C**. However, we observed that the GLM models also perform well in this validation (**Fig. S5**). Hence, considering the opportunity of using a pre-trained model, it might be beneficial to use the GLM, as it was slightly more accurate, and it can be applied fast. In comparison to the LSM, that still needs to evaluate an artificial neural network, the GLM simply performs a product of the input vector and the model coefficients, which is a very cheap operation in terms of computational costs.

## Methods

### Dataset and classification scenarios

The data investigated for this study were derived from experiments with awake, head-fixed mice trained to perform a go / no-go whisker stimulus detection task [58]. Mice were water-restricted and learned that they are rewarded by a drop of water when licking within 500 ms after the onset of a 100-ms sinusoidal vibration of a selected whisker. Before mice were trained to perform this task, they were habituated to the head-fixed setting, which was important to enable the recording of neural activity in the cortical column in the barrel cortex associated with the stimulated whisker. This was done with multi-electrode-array (MEA) silicon probes through all cortical layers using 2-shank-64-channel probes with a distance between the shanks of either 200 or 250 μm. (Neuro Nexus, Ann Arbor, United States, or Cambridge Neurotech, Cambridge, United Kingdom, respectively). Each shank has 32 electrodes and the distance between electrodes is 25 μm, which enables the observation of neural activity over an extent of ∼0.8 mm.

For more details about the mouse line, behavioral task training, habituation, surgery, MEA recordings and the layer-electrode association using current source density (CSD) plots we refer to Vandevelde *et al.* [58]. After inspection of the CSD plots, we decided to use one electrode per layer to derive layer-specific LFPs from the MEA recordings. Psychometric curves (**Fig. 1C**) differ slightly from those shown in our previous study [58], because we excluded trials with a lick response within the first 100 ms after stimulus onset as the muscle movement related to licking caused artifacts in the recordings. For more information on the preprocessing of the LFP traces and LFP features we refer to **SI Materials and Methods, Section 1**.

### Machine learning grid search and cross-validation

As it is not possible to know a priori which algorithm is able to train the most accurate model on a given classification scenario, we performed a grid search including the aforementioned algorithms. For each algorithm, a set of different parameter settings was used to find the algorithm setting leading to the best performance, which also represents the amount of information an algorithm can extract from the dataset (see **SI Materials and Methods, Section 2, Table T1** for details about the parameter grid, software packages used and hardware specifications). Hence, each classifier setting was applied on the data to train a model and evaluated on different classification scenarios specified by the feature vector used and if it was applied for stimulus detection or response prediction. The evaluation of the models, including their generalizability, was done using a training set for training the model, a validation set used to determine the algorithm-parameter-setting achieving the highest validation accuracy and finally evaluated on a testing set composed of samples neither used for training nor for selecting the best parameter setting. This cross-validation concept was repeated with ten different random but stratified splits, and the training or testing accuracy achieved by one model was averaged over the splits. Stratification ensures that the label-balance, proportion of positive labels in the dataset, remains exactly 50% in training, validation and testing set. Given a certain classification scenario, 20% of the samples were used for testing. The remaining 80% of the data were used for training and validation, again using a split of 80% and 20%, respectively.

The feature vectors used by the classifiers are specified by (i) the type RAW or FFT, (ii) the time window pre, peri, or both and (iii) the cortical layer of the recording electrode from which the LFP was derived, L2/3, L4, L5 and L6. Regarding the layer specification, two more variants were included to assess if we lose information by using the averaged signal from all layers (AVG) and if we gain information by concatenating feature vectors of all layers (ALL). To investigate the capability of classifiers to detect stimuli from different stimulus intensities, a comparison was done between scenarios in which all trials from all intensities were included (ALL intensities) and scenarios in which only clear stimulus trials (catch trials and 100% intensity trials) were included. The labeling of the trials was done, based on the presence (intensity >0%) or absence (catch trials) of a stimulus, considering the SD model training. For the RP scenarios, the positively labeled trials are those with a lick detected within the time +100 ms to 500 ms relative to stimulus onset. In all classification scenarios, we used perfectly balanced datasets, which means that each subset contains the same number of positive trials (stimulus trial or response trial) and negative trials (catch trial or no response). This was achieved by down-sampling trials from the overrepresented label and due to the expected 50% positives in the data, we can assume that the baseline accuracy achieved by random predictions will converge to 50%.

### Application cases and leave-one-mouse-out (LOMO) validation

For the stimulus detection (SD) cases, the ML labels differentiate between catch-trials and stimulus trials, considering the different intensities distinctively. Hence, one expects to have a clear separation of the trials for high intensity, while the detection of stimulus trials with 20% intensity is expected to be most difficult for the classifier. For the intensity classification (IC) cases, the labels differentiate the trials by a certain pair of intensities. Again, similar to the initial grid search, the scenarios were balanced with respect to the labeling by down-sampling trials from the overrepresented label. The same cross-validation concept was applied as it was done during the grid search, and the reported accuracies are those achieved on the testing sets. For the leave-one-mouse-out (LOMO) validation, we first separated the data in order to have all trials from one mouse in the testing set and the trials from the other three mice in the training set. Then both sets, training and testing, were balanced according to the labels. To have a more reliable estimation for the model accuracy, we created bootstrap samples from the training data used to train ten randomly different models. These models were then evaluated based on the mouse-specific testing set, and the corresponding accuracy values are reported in the box plots.

The LSM was not included in the grid search due to limitations regarding its installation on the HPC system, however, a smaller grid search could be applied on a local machine to perform a comparatively fair hyperparameter tuning also for the LSM. To provide a more comprehensive view on the runtime comparison, we decided to include the *slim GLM*, a model which applies the same settings as the *best GLM* but with a lower number of maximum iterations, using the default value of 100. This allows the algorithm to terminate the optimization procedure earlier, resulting in lower training time which does not necessarily result in lower predictive performance.

## Acknowledgements

We gratefully acknowledge the funding from the Carl-Zeiss Foundation (0563-2.8/738/2) initiative “Emergent Algorithmic Intelligence.” This work was supported by grants from the Deutsche Forschungsgemeinschaft (LU375/11-1 to H.J.L. and STU544/3-1 and STU544/4-1 to M.C.S.).

Parts of this research were conducted using the supercomputer MOGON 2 and/or advisory services offered by Johannes Gutenberg-University Mainz (hpc.uni-mainz.de), which is a member of the AHRP (Alliance for High Performance Computing in Rhineland-Palatinate, www.ahrp.info) and the Gauss Alliance e.V. The authors gratefully acknowledge the computing time granted on the supercomputer MOGON 2 at Johannes Gutenberg-University Mainz (hpc.uni-mainz.de).

## Author Contributions

**SA** Conceptualization, Data curation, Data preprocessing, Formal analysis, Investigation, Software, Visualization, Writing article **JRV** Performing behavioral experiments and recordings, Data acquisition, Pre-analysis, Data curation, Data preprocessing **EV** Software, Data preprocessing, Visualization, Editing article **GB** Software, Editing article **DB** Visualization, Editing article **MCS** Conceptualization, Data curation, Investigation, Editing article **HJL** Conceptualization, Investigation, Editing article **IH** Conceptualization, Data preprocessing, Software, Formal analysis, Investigation, Editing article

## Supporting information captions

Figure S1 – Comparison between “case best” model and “generic best” model

Figure S2 – Runtime comparison between the classifiers

Figure S3 – Random Forest feature weights of PERI-stimulus FFT features

Figure S4 – Distribution of falsely predicted trials

Figure S5 – Generalizability analysis with GLMs SI Materials and Methods

– **Section 1 – LFP preprocessing and LFP features**
– **Section 2 – Grid search parameters and details about soft- and hardware used**
  - **Table T1 – Grid search parameter settings**
– **Section 3 – Soft- and hardware specifications**

**SI References**

## References

1. Rapeaux AB, Constandinou TG. Implantable brain machine interfaces: first-in-human studies, technology challenges and trends. Current opinion in biotechnology. 2021;72:102–11.

2. Sitaram R, Ros T, Stoeckel L, Haller S, Scharnowski F, Lewis-Peacock J, et al. Closed-loop brain training: the science of neurofeedback. Nat Rev Neurosci. 2017;18(2):86–100.

3. Semprini M, Laffranchi M, Sanguineti V, Avanzino L, De Icco R, De Michieli L, et al. Technological approaches for neurorehabilitation: from robotic devices to brain stimulation and beyond. Frontiers in neurology. 2018;9:212.

4. Rezeika A, Benda M, Stawicki P, Gembler F, Saboor A, Volosyak I. Brain–computer interface spellers: A review. Brain sciences. 2018;8(4):57.

5. Sunny T, Aparna T, Neethu P, Venkateswaran J, Vishnupriya V, Vyas P. Robotic arm with brain– computer interfacing. Procedia Technology. 2016;24:1089–96.

6. Krucoff MO, Rahimpour S, Slutzky MW, Edgerton VR, Turner DA. Enhancing nervous system recovery through neurobiologics, neural interface training, and neurorehabilitation. Frontiers in neuroscience. 2016;10:584.

7. Guggenmos DJ, Azin M, Barbay S, Mahnken JD, Dunham C, Mohseni P, et al. Restoration of function after brain damage using a neural prosthesis. Proceedings of the National Academy of Sciences. 2013;110(52):21177–82.

8. Petschenig H, Bisio M, Maschietto M, Leparulo A, Legenstein R, Vassanelli S. Classification of Whisker Deflections From Evoked Responses in the Somatosensory Barrel Cortex With Spiking Neural Networks. Frontiers in neuroscience. 2022;16.

9. Buzsáki G, Anastassiou CA, Koch C. The origin of extracellular fields and currents—EEG, ECoG, LFP and spikes. Nature reviews neuroscience. 2012;13(6):407–20.

10. Rossant C, Kadir SN, Goodman DF, Schulman J, Hunter ML, Saleem AB, et al. Spike sorting for large, dense electrode arrays. Nature neuroscience. 2016;19(4):634–41.

11. Feldmeyer D, Brecht M, Helmchen F, Petersen CCH, Poulet JFA, Staiger JF, et al. Barrel cortex function. Prog Neurobiol,. 2013;103:3–27.

12. Staiger JF, Petersen CCH. Neuronal Circuits in Barrel Cortex for Whisker Sensory Perception. Physiol Rev. 2021;101(1):353–415. doi: 10.1152/physrev.00019.2019.

13. Wang X, Magno M, Cavigelli L, Mahmud M, Cecchetto C, Vassanelli S, et al., editors. Rat cortical layers classification extracting evoked local field potential images with implanted multi-electrode sensor. 2018 IEEE 20th International Conference on e-Health Networking, Applications and Services (Healthcom); 2018: IEEE.

14. Wang X, Magno M, Cavigelli L, Mahmud M, Cecchetto C, Vassanelli S, et al., editors. Embedded classification of local field potentials recorded from rat barrel cortex with implanted multi-electrode array. 2018 IEEE Biomedical Circuits and Systems Conference (BioCAS); 2018: IEEE.

15. Buccelli S, Bornat Y, Colombi I, Ambroise M, Martines L, Pasquale V, et al. A neuromorphic prosthesis to restore communication in neuronal networks. IScience. 2019;19:402–14.

16. Werner T, Vianello E, Bichler O, Garbin D, Cattaert D, Yvert B, et al. Spiking neural networks based on OxRAM synapses for real-time unsupervised spike sorting. Frontiers in neuroscience. 2016;10:474.

17. Steriade M, Nunez A, Amzica F. A novel slow (< 1 Hz) oscillation of neocortical neurons in vivo: depolarizing and hyperpolarizing components. Journal of neuroscience. 1993;13(8):3252–65.

18. Stüttgen MC, Ruter J, Schwarz C. Two psychophysical channels of whisker deflection in rats align with two neuronal classes of primary afferents. J Neurosci. 2006;26(30):7933–41.

19. Stüttgen MC, Schwarz C. Psychophysical and neurometric detection performance under stimulus uncertainty. Nat Neurosci. 2008;11:1091–9.

20. Stüttgen MC, Schwarz C. Integration of vibrotactile signals for whisker-related perception in rats is governed by short time constants: comparison of neurometric and psychometric detection performance. J Neurosci. 2010;30(6):2060–9.

21. Andersen RA, Musallam S, Pesaran B. Selecting the signals for a brain–machine interface. Current opinion in neurobiology. 2004;14(6):720–6.

22. Markowitz DA, Wong YT, Gray CM, Pesaran B. Optimizing the decoding of movement goals from local field potentials in macaque cortex. Journal of Neuroscience. 2011;31(50):18412–22.

23. Brigham EO. The fast Fourier transform and its applications: Prentice-Hall, Inc.; 1988.

24. Sederberg AJ, Pala A, Zheng HJV, He BJ, Stanley GB. State-aware detection of sensory stimuli in the cortex of the awake mouse. PLoS Comput Biol. 2019;15(5):e1006716. Epub 20190531. doi: 10.1371/journal.pcbi.1006716. PubMed PMID: 31150385; PubMed Central PMCID: PMCPMC6561583.

25. Breiman L. Random forests. Machine learning. 2001;45(1):5–32.

26. Hochreiter S, Schmidhuber J. Long short-term memory. Neural computation. 1997;9(8):1735–80.

27. Kleinbaum DG, Dietz K, Gail M, Klein M, Klein M. Logistic regression: Springer; 2002.

28. Vecchi E, Pospíšil L, Albrecht S, O’Kane TJ, Horenko I. eSPA+: Scalable Entropy-Optimal Machine Learning Classification for Small Data Problems. Neural Computation. 2022;34(5):1220–55.

29. Horenko I. On a scalable entropic breaching of the overfitting barrier for small data problems in machine learning. Neural Computation. 2020;32(8):1563–79.

30. Maass W. Liquid state machines: motivation, theory, and applications. Computability in context: computation and logic in the real world. 2011:275–96.

31. Stüttgen MC, Rüter J, Schwarz C. Two psychophysical channels of whisker deflection in rats align with two neuronal classes of primary afferents. Journal of Neuroscience. 2006;26(30):7933–41.

32. Temereanca S, Simons DJ. Local field potentials and the encoding of whisker deflections by population firing synchrony in thalamic barreloids. Journal of neurophysiology. 2003;89(4):2137–45.

33. Pinto DJ, Brumberg JC, Simons DJ. Circuit dynamics and coding strategies in rodent somatosensory cortex. Journal of neurophysiology. 2000;83(3):1158–66.

34. Reyes-Puerta V, Sun JJ, Kim S, Kilb W, Luhmann HJ. Laminar and columnar structure of sensory-evoked multineuronal spike sequences in adult rat barrel cortex in vivo. Cerebral Cortex. 2015;25:2001–21.

35. Roy NC, Bessaih T, Contreras D. Comprehensive mapping of whisker-evoked responses reveals broad, sharply tuned thalamocortical input to layer 4 of barrel cortex. Journal of Neurophysiology. 2011;105(5):2421–37.

36. Burnham KP, Anderson DR. Model Selection and Multimodel Inference. Second edition ed: Springer; 2002.

37. Verstraeten D, Schrauwen B, Stroobandt D, Van Campenhout J. Isolated word recognition with the liquid state machine: a case study. Information Processing Letters. 2005;95(6):521–8.

38. Maass W, Natschläger T, Markram H. Real-time computing without stable states: A new framework for neural computation based on perturbations. Neural computation. 2002;14(11):2531–60.

39. Nicolelis MAL. Brain-machine-brain interfaces as the foundation for the next generation of neuroprostheses. Natl Sci Rev. 2022;9(10):nwab206. Epub 20211124. doi: 10.1093/nsr/nwab206. PubMed PMID: 36196121; PubMed Central PMCID: PMCPMC9522427.

40. Lebedev MA, Nicolelis MA. Brain-Machine Interfaces: From Basic Science to Neuroprostheses and Neurorehabilitation. Physiol Rev. 2017;97(2):767–837. doi: 10.1152/physrev.00027.2016. PubMed PMID: 28275048.

41. Rasheed S. A Review of the Role of Machine Learning Techniques towards Brain-Computer Interface Applications. Mach Learn Know Extr. 2021;3(4):835–62. doi: 10.3390/make3040042. PubMed PMID: WOS:000744029500001.

42. Wang XY, Magno M, Cavigelli L, Mahmud M, Cecchetto C, Vassanelli S, et al. Rat Cortical Layers Classification extracting Evoked Local Field Potential Images with Implanted Multi-Electrode Sensor. 2018 Ieee 20th International Conference on E-Health Networking, Applications and Services (Healthcom). 2018. PubMed PMID: WOS:000517603800006.

43. Wang XY, Magno M, Cavigelli L, Mahmud M, Cecchetto C, Vassanelli S, et al. Embedded Classification of Local Field Potentials Recorded from Rat Barrel Cortex with Implanted Multi-Electrode Array. Biomed Circ Syst C. 2018:511–4. PubMed PMID: WOS:000458897900134.

44. Petschenig H, Bisio M, Maschietto M, Leparulo A, Legenstein R, Vassanelli S. Classification of Whisker Deflections From Evoked Responses in the Somatosensory Barrel Cortex With Spiking Neural Networks. Front Neurosci. 2022;16:838054. Epub 20220414. doi: 10.3389/fnins.2022.838054. PubMed PMID: 35495034; PubMed Central PMCID: PMCPMC9047904.

45. Monesi MJ, Accou B, Montoya-Martinez J, Francart T, Van Hamme H, editors. An LSTM based architecture to relate speech stimulus to EEG. ICASSP 2020-2020 IEEE International Conference on Acoustics, Speech and Signal Processing (ICASSP); 2020: IEEE.

46. Gerber S, Pospisil L, Navandar M, Horenko I. Low-cost scalable discretization, prediction, and feature selection for complex systems. Science Advances. 2020;6(5):eaaw0961.

47. Moxon KA, Foffani G. Brain-machine interfaces beyond neuroprosthetics. Neuron. 2015;86(1):55–67.

48. Lee SH, Dan Y. Neuromodulation of brain states. Neuron. 2012;76(1):209–22. doi: 10.1016/j.neuron.2012.09.012. PubMed PMID: 23040816; PubMed Central PMCID: PMCPMC3579548.

49. Stüttgen MC, Schwarz C. Barrel cortex: What is it good for? Neuroscience. 2018;368:3–16.

50. Reyes-Puerta V, Kim S, Sun JJ, Imbrosci B, Kilb W, Luhmann HJ. High stimulus-related information in barrel cortex inhibitory interneurons. Plos Computational Biology. 2015;11(6):e1004121.

51. Bonaiuto JJ, Rossiter HE, Meyer SS, Adams N, Little S, Callaghan MF, et al. Non-invasive laminar inference with MEG: Comparison of methods and source inversion algorithms. Neuroimage. 2018;167:372–83.

52. Martinez D, Clément M, Messaoudi B, Gervasoni D, Litaudon P, Buonviso N. Adaptive quantization of local field potentials for wireless implants in freely moving animals: an open-source neural recording device. Journal of neural engineering. 2018;15(2):025001.

53. Won SM, Cai L, Gutruf P, Rogers JA. Wireless and battery-free technologies for neuroengineering. Nature Biomedical Engineering. 2021:1–19.

54. Jackson A, Mavoori J, Fetz EE. Long-term motor cortex plasticity induced by an electronic neural implant. Nature. 2006;444(7115):56–60.

55. O’Doherty JE, Lebedev MA, Ifft PJ, Zhuang KZ, Shokur S, Bleuler H, et al. Active tactile exploration using a brain-machine-brain interface. Nature. 2011;479(7372):228–32.

56. Carlini N, Wagner D, editors. Towards evaluating the robustness of neural networks. 2017 ieee symposium on security and privacy (sp); 2017: Ieee.

57. Rice L, Wong E, Kolter Z, editors. Overfitting in adversarially robust deep learning. International Conference on Machine Learning; 2020: PMLR.

58. Vandevelde JR, Yang J-W, Albrecht S, Lam H, Kaufmann P, Luhmann HJ, et al. Layer-and cell-type-specific differences in neural activity in mouse barrel cortex during a whisker detection task. Cerebral Cortex. 2022.

59. Chen-Bee CH, Zhou Y, Jacobs NS, Lim B, Frostig RD. Whisker array functional representation in rat barrel cortex: transcendence of one-to-one topography and its underlying mechanism. Frontiers in neural circuits. 2012;6:93.

60. Yamashita T, Vavladeli A, Pala A, Galan K, Crochet S, Petersen SS, et al. Diverse long-range axonal projections of excitatory layer 2/3 neurons in mouse barrel cortex. Frontiers in neuroanatomy. 2018;12:33.

