## Supplementary Material for "Spiking neural networks provide accurate, efficient and robust models for whisker stimulus classification and allow for inter-individual generalization"

Short Title: **Generalizable machine learning models for whisker stimulus classification**

Steffen Albrecht [1,*], Jens R. Vandevelde [1,2], Edoardo Vecchi [3], Gabriele Berra [3], Davide Bassetti [4], Maik C. Stüttgen [5], Heiko J. Luhmann [1,*], Illia Horenko [4,*]

[1] Institute of Physiology, University Medical Center of the Johannes Gutenberg University Mainz, Mainz, Germany

[2] Current address: Center for Integrative Physiology and Molecular Medicine, Saarland University, Homburg, Germany

[3] Institute of Computing, Faculty of Informatics, Universitá della Svizzera Italiana (USI), Lugano, Switzerland

[4] Artificial Intelligence in Mathematics, Department of Mathematics, TU Kaiserslautern, Kaiserslautern, Germany

[5] Institute of Pathophysiology, University Medical Center of the Johannes Gutenberg University Mainz, Mainz, Germany

* corresponding authors

**This PDF file includes:**

SI Results
Figures S1 to S5
SI Materials and Methods
Table T1
SI References

### SI Results

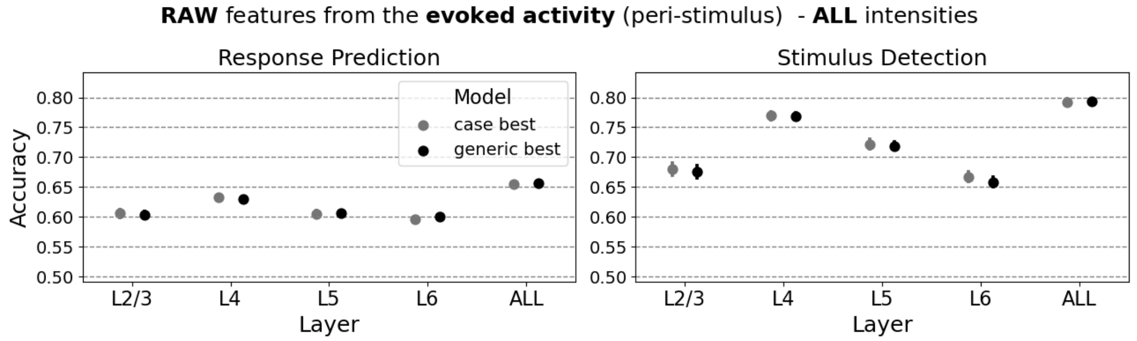
**RAW** features from the **evoked activity** (peri-stimulus) - **ALL** intensities
Response Prediction
Stimulus Detection
Model
case best
generic best
Accuracy
Layer
L2/3
L4
L5
L6
ALL

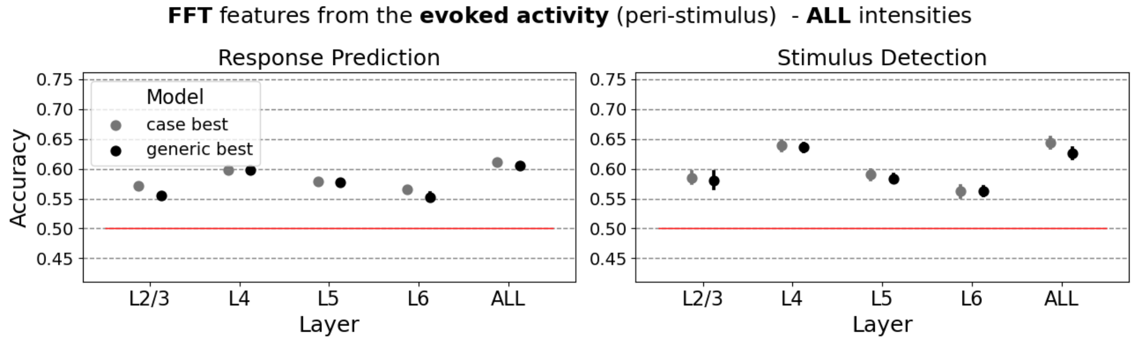
**FFT** features from the **evoked activity** (peri-stimulus) - **ALL** intensities
Response Prediction
Stimulus Detection
Model
case best
generic best
Accuracy
Layer
L2/3
L4
L5
L6
ALL

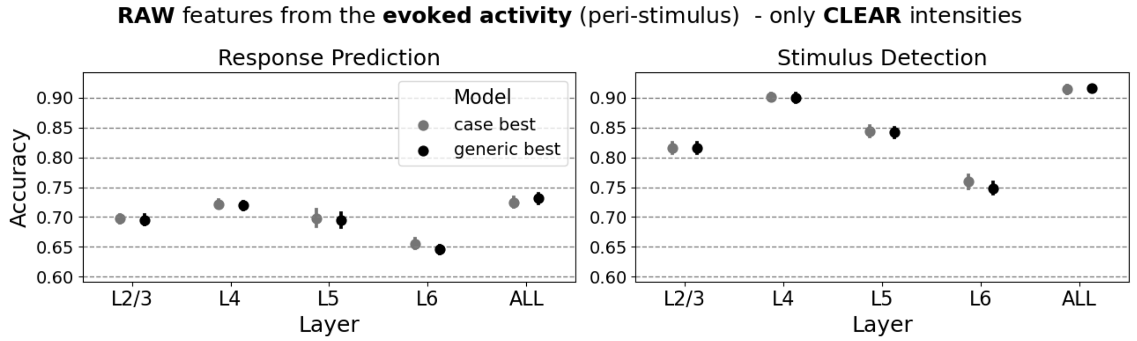
**RAW** features from the **evoked activity** (peri-stimulus) - only **CLEAR** intensities
Response Prediction
Stimulus Detection
Model
case best
generic best
Accuracy
Layer
L2/3
L4
L5
L6
ALL

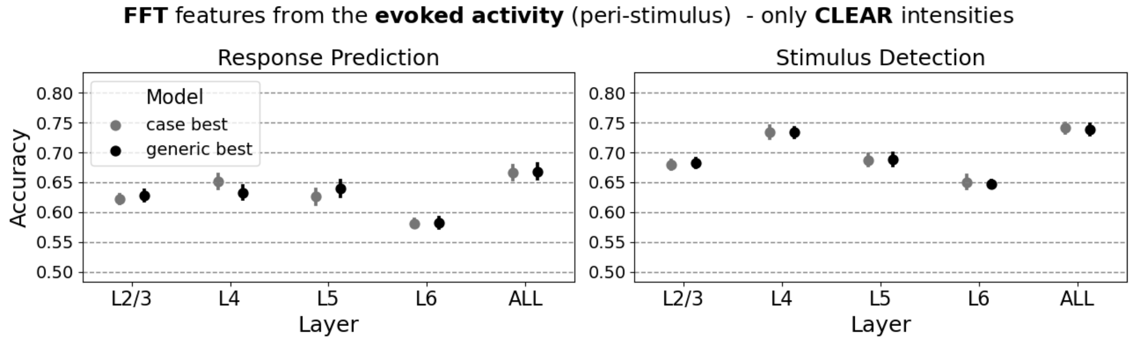
**FFT** features from the **evoked activity** (peri-stimulus) - only **CLEAR** intensities
Response Prediction
Stimulus Detection
Model
case best
generic best
Accuracy
Layer
L2/3
L4
L5
L6
ALL

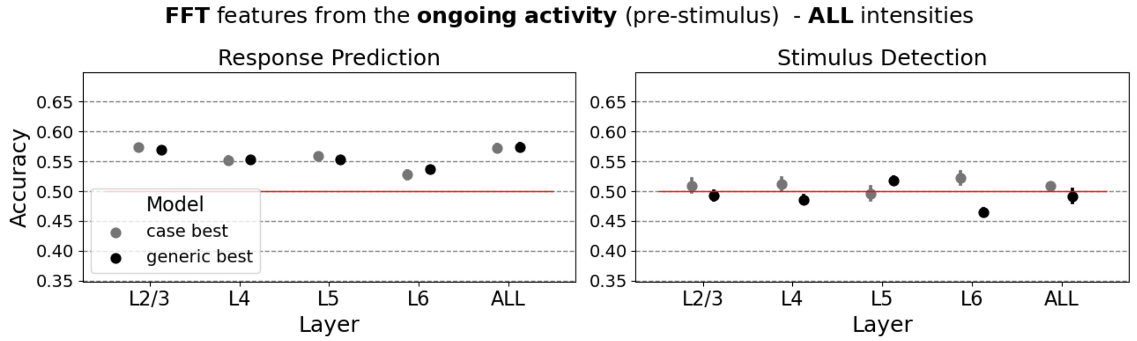
**FFT** features from the **ongoing activity** (pre-stimulus) - **ALL** intensities
Response Prediction
Stimulus Detection
Model
case best
generic best
Accuracy
Layer
L2/3
L4
L5
L6
ALL

**Figure S1 – Comparison between “case best” model and “generic best” model**

Performance described by the testing accuracy from different models applied on the different scenarios as specified by the subplot titles. The grey dots represent the model with the highest validation accuracy for a specific scenario further specified by the cortical layer. The black dots represent the “generic best” model selected based on the highest average accuracy derived from all scenarios for the corresponding feature type. Error bars show the standard error of the mean (SEM). Red lines indicate the accuracy of 50% as expected from an untrained, random guess model. Note that the y-axis scale differs depending on the level of accuracies plotted, however, the range between minimum and maximum y-ticks was set to 0.3 for all subplots.

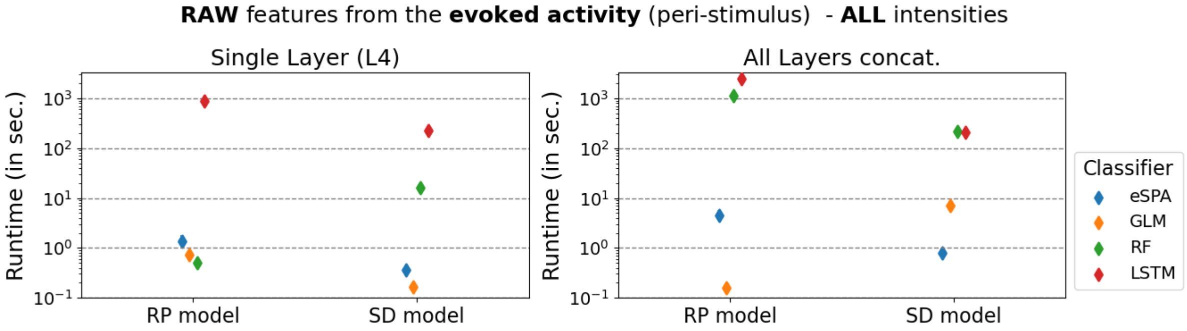
RAW features from the evoked activity (peri-stimulus) - ALL intensities
Single Layer (L4)
All Layers concat.
Runtime (in sec.)
RP model
SD model
Classifier
eSPA
GLM
RF
LSTM

47

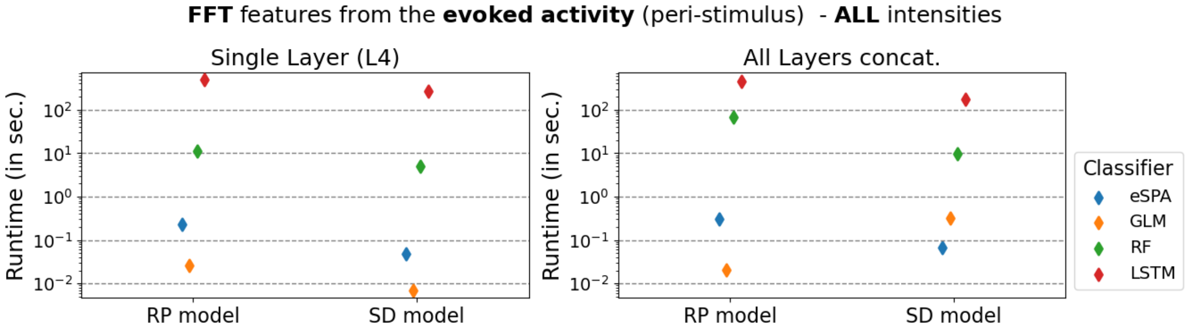
FFT features from the evoked activity (peri-stimulus) - ALL intensities
Single Layer (L4)
All Layers concat.
Runtime (in sec.)
RP model
SD model
Classifier
eSPA
GLM
RF
LSTM

48

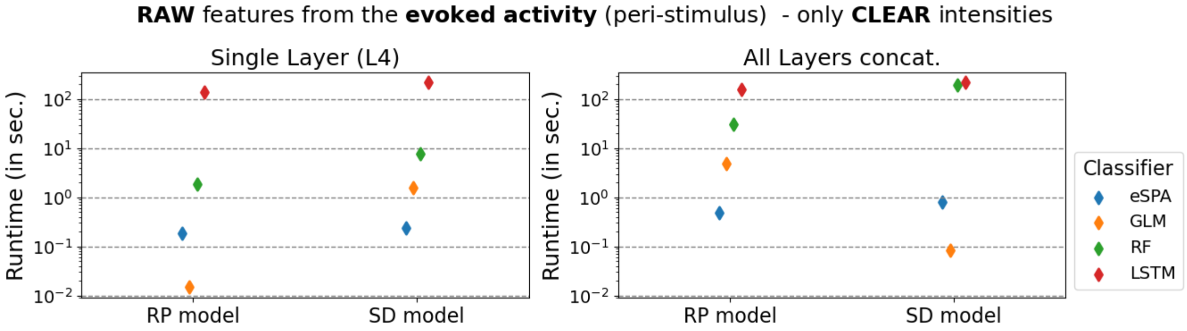
RAW features from the evoked activity (peri-stimulus) - only CLEAR intensities
Single Layer (L4)
All Layers concat.
Runtime (in sec.)
RP model
SD model
Classifier
eSPA
GLM
RF
LSTM

49

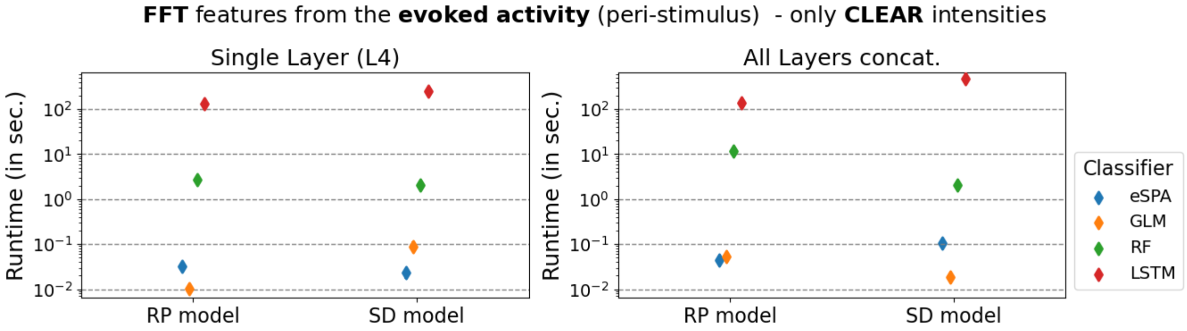
FFT features from the evoked activity (peri-stimulus) - only CLEAR intensities
Single Layer (L4)
All Layers concat.
Runtime (in sec.)
RP model
SD model
Classifier
eSPA
GLM
RF
LSTM

50

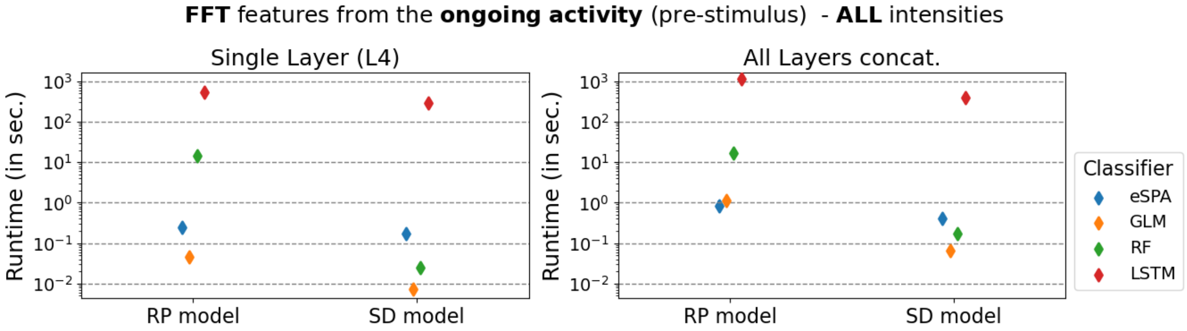
FFT features from the ongoing activity (pre-stimulus) - ALL intensities
Single Layer (L4)
All Layers concat.
Runtime (in sec.)
RP model
SD model
Classifier
eSPA
GLM
RF
LSTM

51

**Figure S2 – Runtime comparison between the classifiers**

Runtime described as the time in seconds a classifier needs to train the model considering the “case best” settings. As the confidence intervals were not visible on this scale, only the mean runtime over the ten cross-validation splits from the grid search are shown.

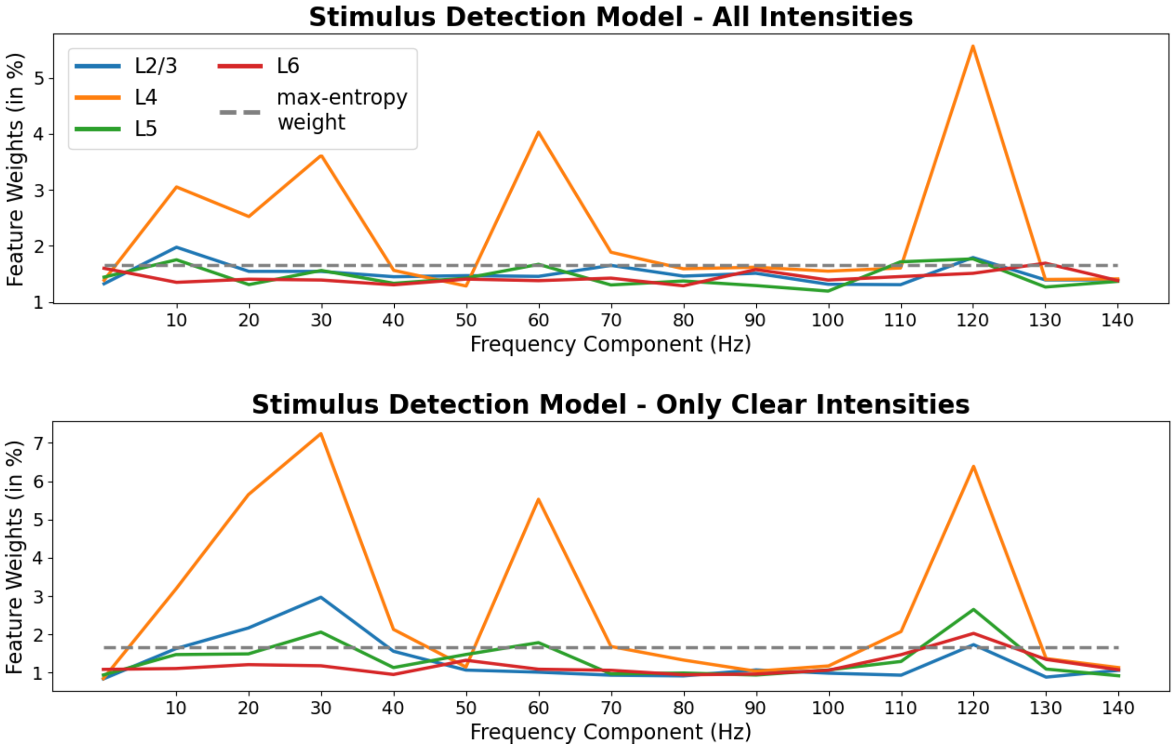


**Figure S3 – Random Forest feature weights of PERI-stimulus FFT features**

Feature weights are represented by the impurity-based feature importance from the random forest algorithm. PERI-stimulus FFT features are available in 10 Hz resolution while the very first feature simply describes the average signal over the given time window. As a reference, the maximum-entropy weight line describes a baseline and represents the uniformly distributed feature weight vector expected from a default and untrained (random) model. The RF model used to derive these feature weights was trained on the concatenated vector including the feature sets from all layers.

#### Section 1 – Distribution of falsely predicted trials

We further investigated the difference between SD and RP models by analyzing the falsely predicted trials over the course of the session. This analysis was motivated by our previous study in which we observed a changing decision criterion related to the decreasing response probability from mice towards the end of the session as mice become more saturated and consequently less motivated to receive the water reward (Vandevelde, Yang et al. 2022). Thus, the mouse emits more false licks in the beginning and less hits in the end of the session. Nevertheless, the response of mice is strongly linked to the presence of the stimulus and our expectation is that the RP models eventually learn to detect the stimulus without being able to extract more information from the signal that is predictive towards the discrimination of successful response and no-response trials. As expected, the RP model generates more False-Negatives in the beginning of the session as the model considers catch trials as no-response trials while mice tried to receive water spontaneously even if there was no whisker stimulus (**Fig. S4**). Towards the end of the session, the RP model positively detects stimulus-trials which are often response trials as well, however, even for strong stimuli mice responded less often in the end of the session resulting in False-Positive predictions of the model. From these observations we conclude that the RP model supposedly predicts the response based on the stimulus detection, without being able to extract statistical patterns predictable towards the behavior.

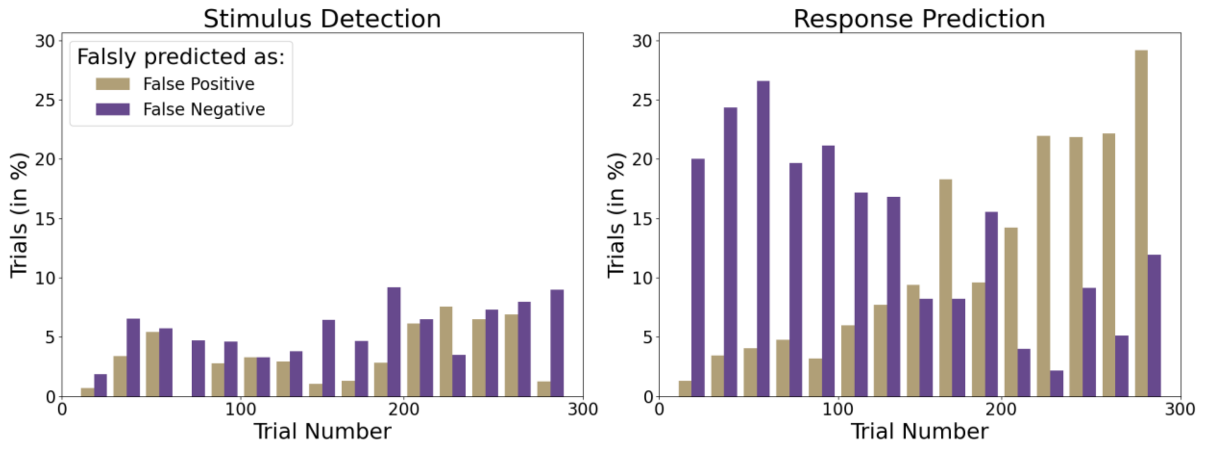


**Figure S4 – Distribution of falsely predicted trials**

Bars represent the proportion of False-Positive and False-Negative predictions from two types of models, the Stimulus Detection model (left) and Response Prediction model (right). Trials were binned over the course of the session with bin width of 20 resulting in 15 bins for 300 trials per session.

#### Section 2 – Additional figure about the generalizability of the GLM models

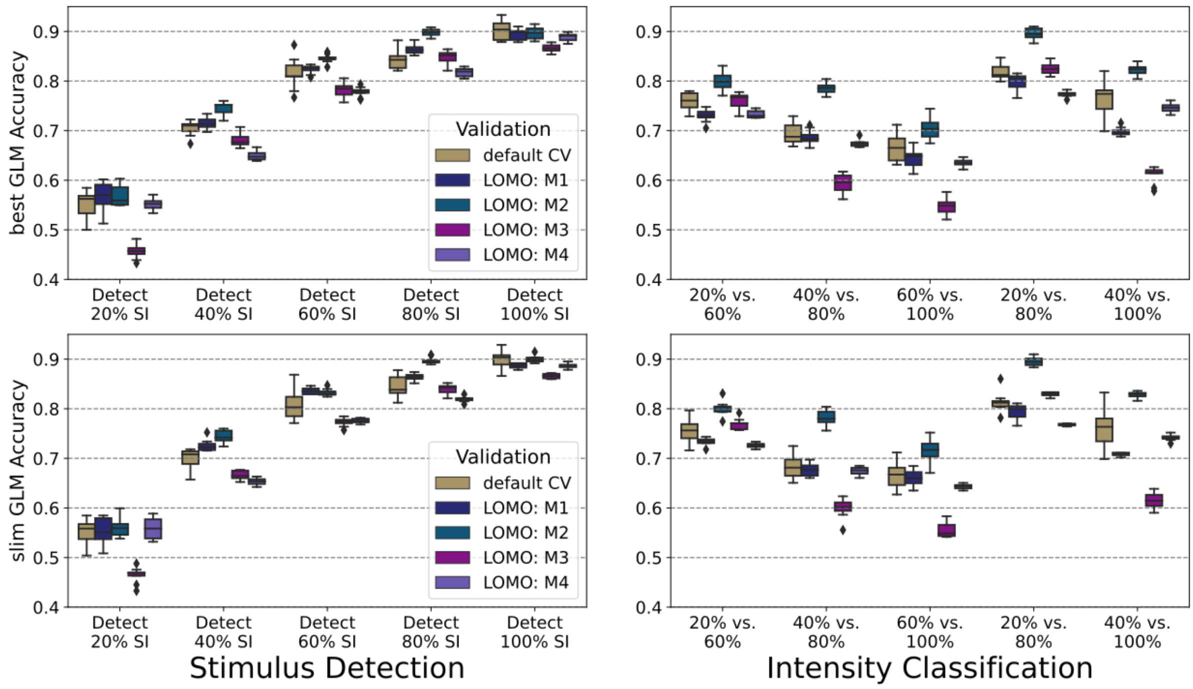


**Figure S5 – Generalizability analysis with GLMs**

As in **Figure 4C**, the additional leave-one-mouse-out (LOMO) is shown for the GLMs used in the final benchmarking. The LOMO validation is compared to the default cross-validation that includes a mixture of trials representing all individuals.

### SI Materials and Methods

#### Section 1 – LFP preprocessing and LFP features

In order to exclude LFP traces contaminated by artifacts, we applied trial-filters for the full range from -420 ms to +120 ms according to the following criteria. Trials in which animals were presumably moving and/or licking randomly had large voltage fluctuations and sometimes burst-like oscillations in activity. Trials in which the signal reached the saturation were identified by applying a cutoff of ±2 mV at any time point within the time range specified above. Bursts were identified by moving time windows of 20 ms in which all electrodes strongly correlate, computed by Pearson's correlation (Schober, Boer et al. 2018). Electric shorts were partly captured by the MEA electrodes resulting in a continuous 50 Hz sinusoidal signal and trials with a strong 50 Hz frequency power were removed. Trials for which a lick was detected within the above-mentioned time window were also excluded.

The RAW features describe the LFP trace at 1000 Hz resolution, resulting in 400 features describing, for instance, the pre-stimulus activity of 400 ms from -400 ms to 0 ms (stimulus onset). Hence, the peri-stimulus activity from 0 to +100 ms is described by a feature vector of length 100 while 500 features represent the full time-window from -400 ms to +100 ms. Using causal Butterworth filtering, the LFP traces were low-pass filtered at 150 Hz and the 47-53 Hz band was removed (Selesnick and Burrus 1998). Using the same time-windows, the Fast-Fourier-Transform (FFT) was applied to derive the strength of the frequency components (Brigham 1988). According to the length of the signal derived from the investigated time windows, the resolution is 2.5 Hz and 10 Hz going up to 147.5 Hz and 140 Hz for the pre- and peri-stimulus activity, respectively. The first FFT feature (0 Hz) represents the constant, which is simply the average over the signal. Eventually, 60 pre-stimulus and 15 peri-stimulus features describe the FFT feature vectors, resulting in 75 features for the full range. Due to the strong differences between ongoing and evoked activity, the FFT was applied to those time-windows separately and the resulting FFT features were merged afterwards in order to create the full (pre- and peri-stimulus) FFT feature vector.

#### Section 2 – Grid search parameters and details about soft- and hardware used

| **Algorithm** | **Parameter Grid** |
|---|---|
| Random Forest (RF) | *n_estimators*: 5, 10, 25, 50, 100, 200, 500, 1000<br>*max_samples*: None, 0.25, 0.5, 0.75<br>*max_features*: None, sqrt, log2, *min_samples_split*: 2, 5, 10<br>*min_samples_leaf*: 1, 2, 5, 10, *criterion*: gini, entropy |
| Generalized Linear Model (GLM) – Logistic Regression | **General parameters**:<br>*max_iter*: 100, 1000, *fit_intersept*: True, False<br>*C*: $1e^{x}$ with $x \in \mathbb{N}$ ranging from -12 to 12 in steps of size 1<br>**Solver settings depending on penalty:**<br>*l1 penalty: liblinear, saga*<br>*l2 penalty: newton-cg, lbfgs, sag, saga*<br>*no penalty: newton-cg, lbfgs, sag, saga* |
| Long Short-Term Memory (LSTM) | *number of hidden units*: 2, 4, 10, 50, 100, 200 |
| Entropy-Optimal Scalable Probabilistic Approximation (eSPA) | *K*: 20, 100<br>*epsilon W*: 0.1, 1.0, 100000.0, 100.0, 1e-05, 1000.0, 10.0, 5e-05, 0.01, 10000.0, 5e-06, 0.0001, 1e-06, 0.001<br>*epsilon CL*: 0.1, 1e-09, 1.0, 100.0, 1e-05, 1e-10, 1e-08, 5e-05, 0.0001, 0.001, 0.01, 1e-12, 0.005, 1e-07, 1e-06, 1e-11, 0.0005<br>*annealing steps*: 40 |
| Liquide State Machine (LSM) | *excitatory neurons*: 100, 250, 500, 1000, 2000<br>*inhibitory neurons*: 10, 25, 100, 250, 500<br>*recurrent neurons*: 50, 250, 500<br>*regularization*: 1e-08, 1e-07, 1e-06, 1e-05, 0.0001, 0.001, 0.01, 0.1, 0.2, 0.5, 1.0, 2.0, 10.0, 100.0 |

**Table T1 – Grid search parameter settings**

#### Section 3 – Soft- and hardware specifications

We used MOGON II the high-performance computing system from the University of Mainz and a local server. We refer to these systems as *mogon* and the *local machine*, respectively. We ran RF and GLM from the sklearn package (Pedregosa, Varoquaux et al. 2011) on mogon. The LSTM was used from the “Deep Learning Toolbox” from MATLAB and eSPA was also performed with MATLAB using own implementations according to the publication of Vecchi *et al.* (Vecchi, Pospíšil et al. 2022). The LSM algorithm was used from a the GitHub repository https://github.com/IGITUGraz/LSM, published by Maass *et al.* (Maass, Natschläger et al. 2002).

The CPUs used on mogon are Intel® Xeon® E5 v4, 2.2GHz and the local machine is equipped with 2 cores, Intel® Xeon® Gold 6240R, 2.4GHz. Due to the required MATLAB licenses the LSTM was ran on the local machine and as there were difficulties with its installation we ran the LSM on the local machine, as well. All other classifiers were used on mogon as we could apply more model training at the same time using the HPC facilities. To make the runtime comparison more reliable, each model was trained using a single thread and the multi thread cores were solely used to train more models in parallel without having a speed up for a single training process. This allowed us to get an approximate runtime comparison as shown in **Figure S2**, with slightly better conditions for the LSTM as the CPUs on the local machine are slightly faster. To reduce the computations on the local machine, the LSM grid search was not applied to all cases shown in **Figure 2**. We focused here on the most important scenarios of training SD models based on all stimulus intensities, but also the CLEAR intensity cases. We considered only the RAW peri-stimulus features here for all L2/3, L4, L5, L5 and ALL, resulting in ten cases in total.

Note that the final runtime comparison (**Fig. 4B**) was completely done on the local machine by retraining also the GLM models using the best setting and the slim setting. Again, only one thread was available for training a single model. In this way we performed an absolutely fair benchmarking as the LSM, best GLM and slim GLM used the same computational resources and CPU parallelization was disabled for the three classifiers.

### SI References


Brigham, E. O. (1988). The fast Fourier transform and its applications, Prentice-Hall, Inc.

Maass, W., T. Natschläger and H. Markram (2002). "Real-time computing without stable states: A new framework for neural computation based on perturbations." Neural computation **14**(11): 2531-2560.

Pedregosa, F., G. Varoquaux, A. Gramfort, V. Michel, B. Thirion, O. Grisel, M. Blondel, P. Prettenhofer, R. Weiss and V. Dubourg (2011). "Scikit-learn: Machine learning in Python." the Journal of machine Learning research **12**: 2825-2830.

Schober, P., C. Boer and L. A. Schwarte (2018). "Correlation coefficients: appropriate use and interpretation." Anesthesia & Analgesia **126**(5): 1763-1768.

Selesnick, I. W. and C. S. Burrus (1998). "Generalized digital Butterworth filter design." IEEE Transactions on signal processing **46**(6): 1688-1694.

Vandevelde, J. R., J.-W. Yang, S. Albrecht, H. Lam, P. Kaufmann, H. J. Luhmann and M. C. Stüttgen (2022). "Layer-and cell-type-specific differences in neural activity in mouse barrel cortex during a whisker detection task." Cerebral Cortex.

Vecchi, E., L. Pospíšil, S. Albrecht, T. J. O'Kane and I. Horenko (2022). "eSPA+: Scalable Entropy-Optimal Machine Learning Classification for Small Data Problems." Neural Computation **34**(5): 1220-1255.
